# Involvement of a serotonin/GLP-1 circuit in adolescent isolation-induced diabetes

**DOI:** 10.1101/2023.06.12.544498

**Authors:** Louis J. Kolling, Kanza Khan, Nagalakshmi Balasubramanian, Deng-Fu Guo, Kamal Rahmouni, Catherine A. Marcinkiewcz

**Author notes:** Corresponding Author: Catherine Marcinkiewcz Department of Neuroscience and Pharmacology, University of Iowa; Iowa City, Iowa, USA.

## Abstract

In 2020, stay-at-home orders were implemented to stem the spread of SARS-CoV-2 worldwide. Social isolation can be particularly harmful to children and adolescents—during the pandemic, the prevalence of obesity increased by ∼37% in persons aged 2-19. Obesity is often comorbid with type 2 diabetes, which was not assessed in this human pandemic cohort. Here, we investigated whether male mice isolated throughout adolescence develop type 2 diabetes in a manner consistent with human obesity-induced diabetes, and explored neural changes that may underlie such an interaction. We find that isolating C57BL/6J mice throughout adolescence is sufficient to induce type 2 diabetes. We observed fasted hyperglycemia, diminished glucose clearance in response to an insulin tolerance test, decreased insulin signaling in skeletal muscle, decreased insulin staining of pancreatic islets, increased nociception, and diminished plasma cortisol levels compared to group-housed control mice. Using Promethion metabolic phenotyping chambers, we observed dysregulation of sleep and eating behaviors, as well as a time-dependent shift in respiratory exchange ratio of the adolescent-isolation mice. We profiled changes in neural gene transcription from several brain areas and found that a neural circuit between serotonin-producing and GLP-1-producing neurons is affected by this isolation paradigm. Overall, spatial transcription data suggest decreased serotonin neuron activity (via decreased GLP-1-mediated excitation) and increased GLP-1 neuron activity (via decreased serotonin-mediated inhibition). This circuit may represent an intersectional target to further investigate the relationship between social isolation and type 2 diabetes, as well as a pharmacologically-relevant circuit to explore the effects of serotonin and GLP-1 receptor agonists.

**Article Highlights:**

- Isolating C57BL/6J mice throughout adolescence is sufficient to induce type 2 diabetes, presenting with fasted hyperglycemia.
- Adolescent-isolation mice have deficits in insulin responsiveness, impaired peripheral insulin signaling, and decreased pancreatic insulin production.
- Transcriptional changes across the brain include the endocannabinoid, serotonin, and GLP-1 neurotransmitters and associated receptors.
- The neural serotonin/GLP-1 circuit may represent an intersectional target to further investigate the relationship between social isolation and type 2 diabetes. Serotonin-producing neurons of adolescent-isolation mice produce fewer transcripts for the GLP-1 receptor, and GLP-1 neurons produce fewer transcripts for the 5-HT_1A_ serotonin receptor.

## Introduction

In 2020, social distancing and stay-at-home orders were implemented as a non-pharmacological measure to stem the spread of SARS-CoV-2 worldwide. This period ranged in duration from 3-30 months (1,2), far greater than the 10-day duration that is known to cause lasting psychiatric effects (3–6). Social isolation can be particularly harmful to children and adolescents (7–9), greatly impacting development and increasing the risk of disease in adulthood (10–14). Short periods of early isolation negatively affect cortical development (15,16), lipid metabolism (17), and decision-making through adulthood (16). Reintegration following isolation does not abrogate these long-term effects (16). During the pandemic, the rate of body weight gain nearly doubled in persons aged 2-19. From August 2019 to August 2020, obesity prevalence increased from 19.3% to 22.4% in this age group (18).

In mice, the age of sexual maturity is often used to define adulthood, which occurs between 8-12 weeks of age (19). Thus, a mouse aged postnatal day 70 (P70) is approximately equivalent to a human aged 20 years (20). In male rats, social isolation throughout adolescence has been shown to impair hepatic insulin sensitivity without affecting fasted blood-glucose levels (21). Adult C57BL/6J (C57) mice isolated for 8 weeks exhibit increased body fat percentage, without an overall change in body weight (22). These effects suggest a possible relationship between social isolation and type 2 diabetes. Relevant neural changes that may bridge social isolation and type 2 diabetes have not been previously assessed, despite the developmental importance of the adolescent period. The activity of serotonin neurons (5-HTNs) is crucial for the regulation of mood and behavior, and adolescent isolation has been reported to decrease the activity of 5- HTNs (23). A subset of 5-HTNs project to the GLP-1-producing preproglucagon neurons (PPGNs) of the nucleus tractus solitarii (NTS), where they differentially modulate the activity of PPGNs by way of 5-HT_1A_ and 5-HT_2C_ receptors (24). A subset of PPGNs project back to the raphe nuclei and modulate the activity of 5-HTNs via the GLP-1 receptor (GLP-1R) (25).

Here, we investigated whether male mice isolated through the end of adolescence (P76) developed type 2 diabetes in a manner consistent with human obesity-induced diabetes. We assessed glucose and insulin tolerance, peripheral and hepatic insulin signaling, pancreatic function, behavior, metabolism, neural gene expression across several brain regions implicated in glucose metabolism, and the transcription levels of receptor genes involved in the 5-HT/GLP-1 circuit.

## Research Design and Methods

### Ethical approval

All animal procedures were reviewed and approved by the UI Office of Animal Resources (Protocol 1032080) that abided by the American Veterinary Medical Association and the National Institutes of Health.

### Animals

Male C57BL/6J mice (catalog #000664, Jackson Labs, Bar Harbor, ME, USA; RRID:IMSR_JAX:000664) were purchased for arrival at P20, shipped five mice per cage. At age P22, half of each cohort (i.e., 10 mice) was moved to individual housing. A total of 80 C57BL/6J male mice were used in our experiments. Female mice were not included in this assessment, because they have been reported to be resistant to the early-adult onset of obesity and diabetes (26–29). This sex difference is thought to be due to the protective effects of estrogen on pancreatic beta cells (30,31).

All mice were housed in a temperature- and humidity-controlled AALAC-approved vivarium at the University of Iowa on a standard 12 h/12 h dark/light (reverse) cycle and in accordance with institutional requirements for animal care. Mice were individually- or group-housed in conventional-style cob-bedding rodent cages with nestlets, containing separate food and water that could be obtained *ad libitum*. Mice were maintained on chow that was composed of 14% kcal fat, 60% kcal carbohydrate, and 26% kcal protein (Catalog #5P76, Land O’Lakes, Arden Hills, MN, USA).

### Behavioral Assessments

All behavior tests took place after roughly 8 weeks of isolation or group housing (**Figure 1A**). Mice were given at least one “rest day” between behavior tests.

**Figure 1.**
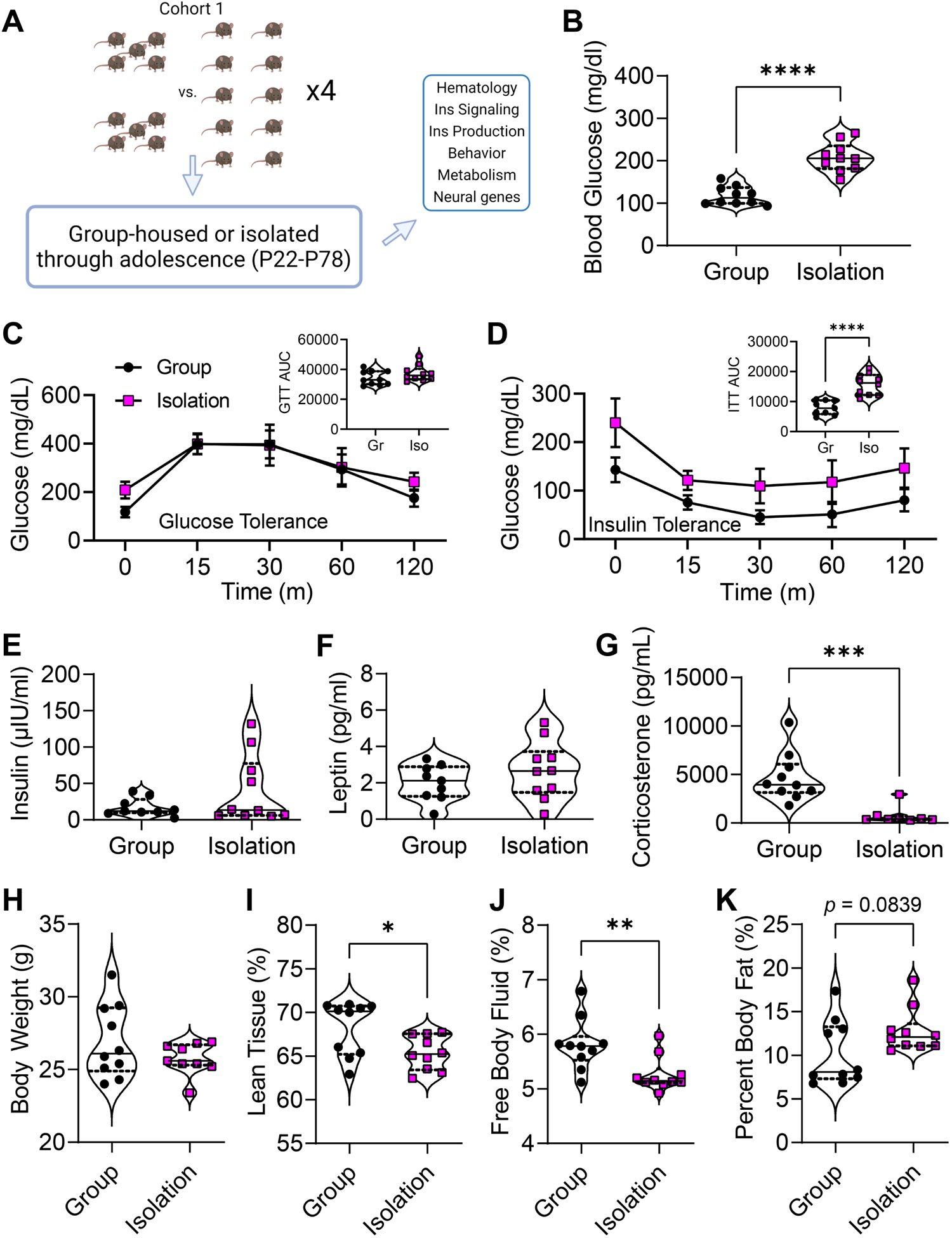
***A:*** Mice within each of the four cohorts were assigned to a housing condition, isolated for 8 weeks, then experimentally assessed (panel created with BioRender.com). ***B:*** Comparison of fasted blood glucose levels, Group = group-housed mice (black), Isolation = adolescent- isolation mice (magenta). ***C:*** Blood glucose levels in response to intraperitoneal injection of glucose, with integrated area under the curve (AUC, ***inset***). ***D:*** Same as ***C***, but in response to intraperitoneal injection of insulin. ***E-G:*** Plasma levels of insulin (***E***), leptin (***F***), and corticosterone (***G***). ***H:*** Comparison of body weights. ***I-K:*** Comparison of percent lean tissue (***I***), free body fluid (***J***), and percent body fat (***K***) as assessed by EchoMRI. \**p* < 0.05, \*\**p* < 0.01, \*\*\**p* < 0.001, \*\*\*\**p* < 0.0001

#### Von Frey

The von Frey test was performed as previously described (32). Briefly, mice were placed in a clear acrylic box (10 x 10 x 15 cm) and placed on the test apparatus. The test apparatus consisted of a raised wire mech platform (holes were 5 x 5 mm). Mice were allowed to acclimate to the acrylic box and test apparatus for three hours on two consecutive days. Mechanical sensitivity was evaluated by applying von Frey filaments (Stoelting, Wood Dale, IL, USA) of varying strengths to the plantar surface of each hind paw. The number of responses out of five applications, per filament per paw, was recorded and used to calculate the 50% withdrawal threshold.

#### Hargreaves

Mice were allowed to acclimate in a clear acrylic box (10 x 10 x 15 cm) on the test apparatus for at least three hours on two consecutive days prior to testing. The test apparatus was an IITC Life Science heated base (Model 400) with temperature maintained at 30° C. Thermal sensitivity was evaluated by focusing a heat-generating light beam onto the plantar surface of each hind paw. The time required to evoke a paw withdrawal was recorded three times per paw and averaged to calculate the paw withdrawal latency.

#### Social interaction test

The social interaction test was performed as previously described (32). Briefly, the animal was placed in the central chamber of a three-chamber arena (20 lux) and allowed to explore the environment for 10 min. Following this, a novel C57BL/6J mouse (“stranger mouse”) of the same sex and relative age (+/- 2 weeks) as the test mouse was placed into one of the side chambers under a metal enclosure. An empty metal enclosure was placed in the alternate side chamber. The test mouse was allowed to explore the environment and interact with the stranger mouse for 10 min. The location of the stranger mouse was alternated between the right and left chambers to control for any potential side preferences. Behavior was recorded by an overhead camera and scored by a researcher blinded to experimental groups. The total time spent interacting with the stranger mouse, and with the novel object (empty enclosure) was analyzed over a 10 min period for each trial, three trials per mouse in total.

#### Body composition analysis

Body composition was assessed *in vivo* using an EchoMRI machine (EchoMRI LLC; RRID:SCR_017104) and manufacturer software. After calibration of the machine, mice were gently restrained within a 2” diameter sample tube. The sample tube was inserted into the EchoMRI, and mice were assessed for fat mass, lean mass, total water, and free water. Body fat percentage was calculated as: bf% = ((fat mass (g))/(lean mass (g) – free water (g))) × 100% (33).

### Hematology

Glucose tolerance tests were performed after an 18-hour fast starting from the beginning of the dark phase (34). After initial determination of fasting blood glucose, 2 g glucose per kg of body weight was administered by intraperitoneal injection from a 10% glucose solution in PBS (University of Virginia Vivarium Protocols, Susanna R. Keller). Glucose levels were measured from collected tail blood samples taken at 0 min (before injection), 15, 30, 60, and 120 min using a OneTouch Ultra 2 blood glucose meter (LifeScan, Malvern, PA, USA). Integrated area under the curve (AUC) was computed to determine clearance.

Intraperitoneal insulin tolerance tests were performed after a 5-hour fast starting from the beginning of the light phase. A stock solution of Humulin R (Eli Lilly) was diluted to a concentration of 2 mIU/ml and intraperitoneally injected at a dose of 1 unit per kg of body weight. Blood glucose levels were measured from tail blood samples taken at 0 min (before injection), 15, 30, 60 and 120 min using a OneTouch Ultra 2 blood glucose meter. AUC was computed to determine clearance as above.

### Biochemistry

#### Insulin signaling assay

Mice were fasted overnight, then anesthetized with intraperitoneal injection of a ketamine-xylazine cocktail before intravenous injection of either vehicle (PBS) or insulin (2.5 units/kg body weight). 15 min following injection, insulin-sensitive tissues (liver, white adipose tissue, brown adipose tissue, and soleus muscle) were harvested from each mouse. Proteins were extracted by homogenizing tissues in tissue lysate buffer (50 mM HEPES, pH 7.5, 150 mM NaCl, 1 mM MgCl_2_, 1 mM CaCl_2_, 10 mM NaF, 5 mM EDTA, 1% Triton, 2 mM sodium orthovanadate and Roche cocktail protease inhibitor tablet). Protein samples (20 µg) were subjected to SDS PAGE, electro-transferred on a polyvinylidene fluoride membrane, then probed with primary rabbit antibodies (1:1,000) targeting AKT (Cell signaling, cat.#: 9272, RRID:AB_329827) and p-AKT(Ser473) (Cell Signaling, cat.#: 4060, RRID:AB_2315049). Membranes were subsequently probed with secondary anti-rabbit antibody (1:10,000). As a loading control, we used β-actin which was targeted with primary mouse antibody (1:25,000, Proteintech, cat#: 60008-1-Ig) followed with anti-mouse secondary antibody (1:10,000). Protein expression was visualized with an Amersham ECL detection kit (Cytiva).

#### ELISA

Plasma preparations were performed using a heparin-free method (University of California, San Diego Metabolomics Protocols and Workflows, Sierra Simpson). Plate-based sandwich ELISAs were performed for insulin (1:4 dilution, RayBiotech, cat# ELM-Insulin-1), leptin (1:40 dilution, RayBiotech, cat# ELM-Leptin-1) and corticosterone (1:1 dilution, Enzo Life Sciences, cat# ADI-900-097) using manufacturer protocols. Samples were run in duplicate, and plates were analyzed using a BioTek Cytation 5 Imaging Reader (RRID:SCR_019732).

### Immunohistochemistry

Pancreata were embedded in paraffin, sectioned at 5 µm, stained with hematoxylin, and chromogenically stained for either insulin or glucagon. Images of pancreata were acquired using an Olympus VS200 slide scanner in the brightfield configuration at 20X. Images were analyzed using FIJI software, based on an approach adapted from Poudel 2016 (35). Islets were morphologically and histologically identified as encapsulated areas devoid of hematoxylin staining and blood cells. ROIs were manually drawn around all islets in a full-color image, and then masked over a corresponding image rendered in grayscale. The “blue” grayscale channel, devoid of hematoxylin staining, was then thresholded using a consistent value of 106. The total islet area per pancreas and the percentage of islet area that was immunopositive for insulin or glucagon stain was calculated.

### Metabolic Assessment

Mice were metabolically profiled using indirect calorimetry by the University of Iowa Metabolic Phenotyping Core. Mice were housed in Promethion cages (Sable Systems International, North Las Vegas, USA) to determine oxygen consumption (VO2; ml/kg/min), thermogenesis, respiratory exchange ratio (RER; VO2/VCO2), locomotor activity, ingestive behavior (caloric and water consumption), and overall interactive behavior with the cage environment. All data were recorded in 5 min intervals, with each interval measurement representing the average value during a 30 s sampling period per cage. In total, mice were housed in the Promethion system for 4 d. No data were acquired during the first 2 d in the chambers to permit acclimation to the environment, and then experimental data were computed for each light/dark phase over 2 d (36). Time 0 reported for the analyses represents the start of the dark or light cycle, respectively.

### RT-qPCR

Brains were flash-frozen on dry ice and stored at −80° C until dissection of brain regions was performed. Anatomically-distinct brain regions were extracted using the Allen Brain Atlas as a guide. The following brain regions were collected: main olfactory bulb (OB), suprachiasmatic nucleus (SCN), a region containing both the arcuate nucleus and ventromedial hypothalamus (ARC/VMH), the lateral hypothalamic area (LHA), periaqueductal grey (PAG), tissue punches containing the lateral dorsal tegmental nucleus (LDT), and the rostral ventromedial medulla (RVM).

Total RNA from each dissected region was extracted using a TRIzol reagent (Ambion, Life Technologies, USA) as described previously (37). DNAse digestion was then performed using a DNA-free™ DNA Removal Kit (Life Technologies, USA). The purity and quantity of RNA were determined using a NanoDrop 1000 spectrophotometer (Thermo Fisher Scientific, USA). cDNA preparation of total mRNA was performed using the iScript cDNA Synthesis Kit (Bio-Rad Laboratories, CA, USA) according to the manufacturer’s protocol. RT-qPCR for target genes was performed using SYBR green qPCR master mix (Bio-Rad Laboratories, USA) and primers (**Supplementary Table 1**) with a CFX96 Real-time-PCR system (Bio-Rad Laboratories, USA). Fold changes in mRNA levels were determined for each gene after normalizing to β-actin Ct values using the fold change 2^−ΔΔCT^ method (38).

### In-situ hybridization in mouse brain tissue (RNAscope)

Coronal sections (25 µm) containing raphe nuclei or NTS were mounted on histobond glass slides. RNAscope was performed according to the ACDbio protocol. In raphe nuclei, the following probes/channels were used: *Tph2*/C1, *Glp1r*/C3. In NTS: *Gcg*/C1, *Htr1a*/C2, *Htr2c*/C3. Images were acquired on an Olympus VS200 slidescanner at 20X, and images were analyzed using Qu-Path 0.4.3 software. The subcellular detection function was used in tandem with pixel classification and cell detection to determine puncta counts per *Tph2*-positive and *Gcg*-positive cell.

### Experimental design and statistical analyses

Data analyses and statistical tests were performed using Prism version 9 (GraphPad Software, Inc., La Jolla, CA, USA; RRID:SCR_002798). Outliers were first determined using a ROUT test (Q = 1%). Data were then checked for normal distribution and homogeneity of variance using the Fmax test. Where data violated homogeneity of variance, a non-parametric analysis was used. All data are reported as the mean ± SD unless otherwise noted. Statistical difference was defined at the 95% confidence interval (or α ≤ 0.05), and all statistical tests used a two-tailed hypothesis. Exact *p* values, statistical tests, and specific variables required for independent replication are found in the **Supplemental Materials**.

## Results

### Adolescent isolation induces type 2 diabetes and alters body composition

After weaning to group-housed or isolated caging (age P22), mice were maintained for a period of 8 weeks (to P78) before performing behavioral or physiological assessments (**Figure 1A**). In mice, the onset of type 2 diabetes is defined by a fasted blood glucose level above 150 mg/dl (39). Adolescent-isolation mice exceed this threshold and had significantly higher blood glucose levels than group-housed controls (*p* < 0.0001) (**Figure 1B**). While no difference was observed throughout a glucose tolerance test (GTT) (**Figure 1C**), adolescent-isolation mice had significantly higher integrated area under the curve (AUC) throughout an insulin tolerance test (ITT) compared to group-housed mice (*p* < 0.0001) (**Figure 1D**). Using commercially-available ELISA kits, we found no difference in plasma insulin concentrations or plasma leptin concentrations. However, we did observe a stark reduction in plasma corticosterone in adolescent-isolation mice (*p* = 0.0006) (**Figure 1E-G**). We further examined body weight and body composition using EchoMRI. Body weights did not differ significantly between housing conditions (**Figure 1H**). Adolescent-isolation mice had less lean tissue, less free fluid, and showed a trend toward increased body fat (**Figure 1I-K**).

### Adolescent isolation impairs peripheral insulin signaling and pancreatic function

Since the insulin tolerance test suggested a deficit in insulin signaling (**Figure 1**), we next examined insulin signaling at various tissue levels. The ratio of phosphorylated AKT (p-AKT; Protein Kinase B) to unphosphorylated AKT is often used to infer the sensitivity of peripheral tissues to insulin. AKT is phosphorylated downstream of the insulin receptor kinase, and in type 2 diabetes, the phosphorylation of AKT is further inhibited via an inositol kinase pathway (40). We gathered tissue from several insulin-sensitive areas and used western blotting to determine the p-AKT/AKT ratio with and without stimulation with insulin. In liver tissue, we noted no difference in p-AKT/AKT ratio (**Figure 2A**). In muscle tissue, we observed a reduction in the relative p-AKT/AKT ratio in adolescent-isolation mice (*p* < 0.0001) (**Figure 2B**). We observed no difference in insulin signaling within brown adipose tissue (**Figure 2C**) or white adipose tissue (**Figure 2D**).

**Figure 2.**
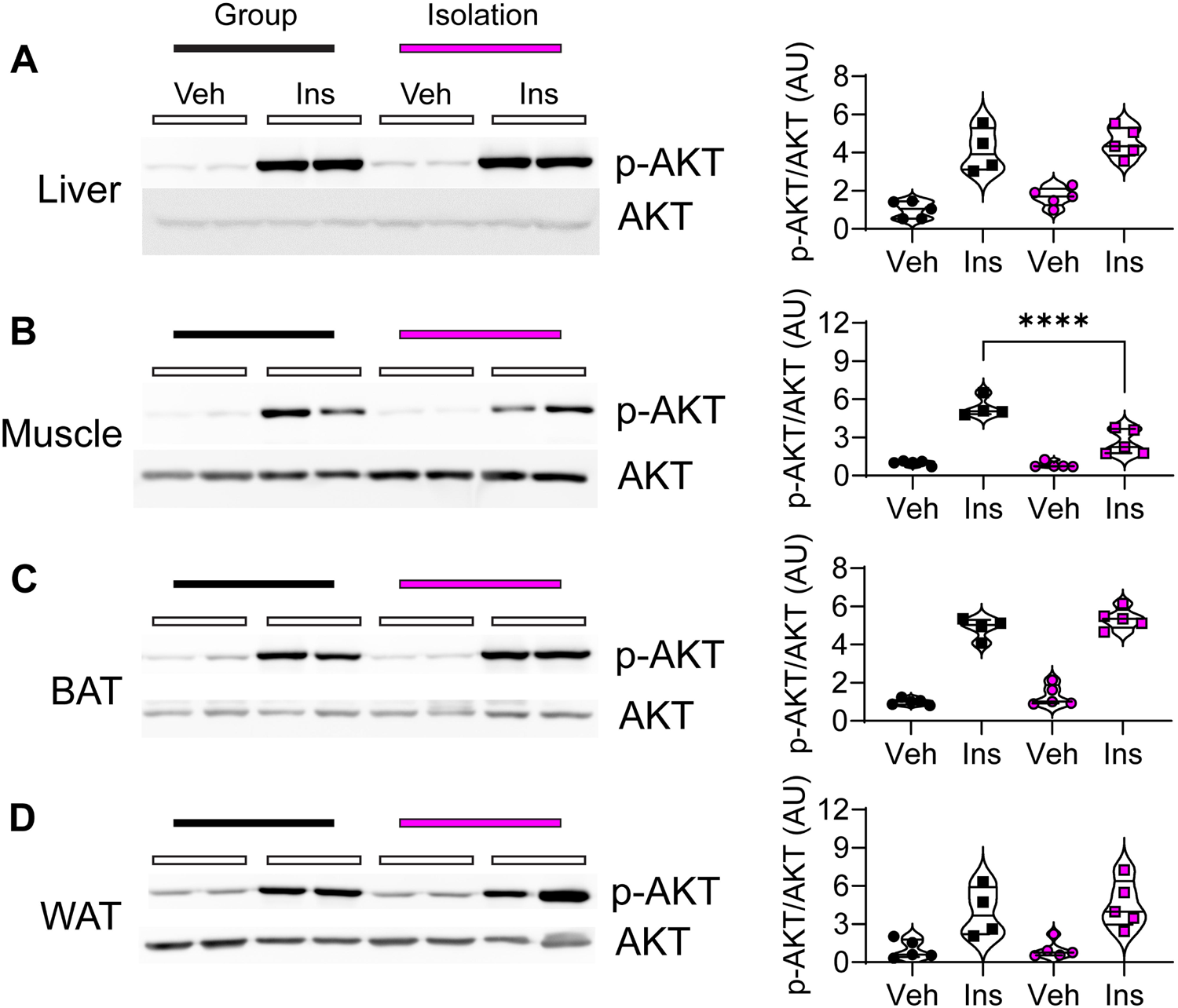
Insulin signaling in various tissue types as determined by AKT signaling. Two animals per treatment (insulin or vehicle injection) from each housing condition (group-housed or adolescent-isolation) were blotted for phosphorylated AKT (p-AKT) or base state AKT (blots, left). Corresponding graphs plot the ratio of p-AKT/AKT as determined using quantitative densitometry (graphs, right). Measurements were performed for liver (***A***), skeletal muscle (***B***), brown adipose tissue (BAT, ***C***) and white adipose tissue (WAT, ***D***). \*\*\*\**p* < 0.0001

Adolescent-isolation mice had an altered ITT without changes in plasma insulin concentrations, so we next harvested and examined pancreata. Formalin-fixed and paraffin-embedded sections were stained with hematoxylin, then immunohistochemistry was performed for insulin or glucagon. Adolescent-isolation mice had less insulin staining as a percentage of total islet area (*p* = 0.0156) (**Figure 3A**), with no change in glucagon staining as a percentage of total islet area (**Figure 3B**). We observed no change in total islet area between groups (**Figure 3C**).

**Figure 3.**
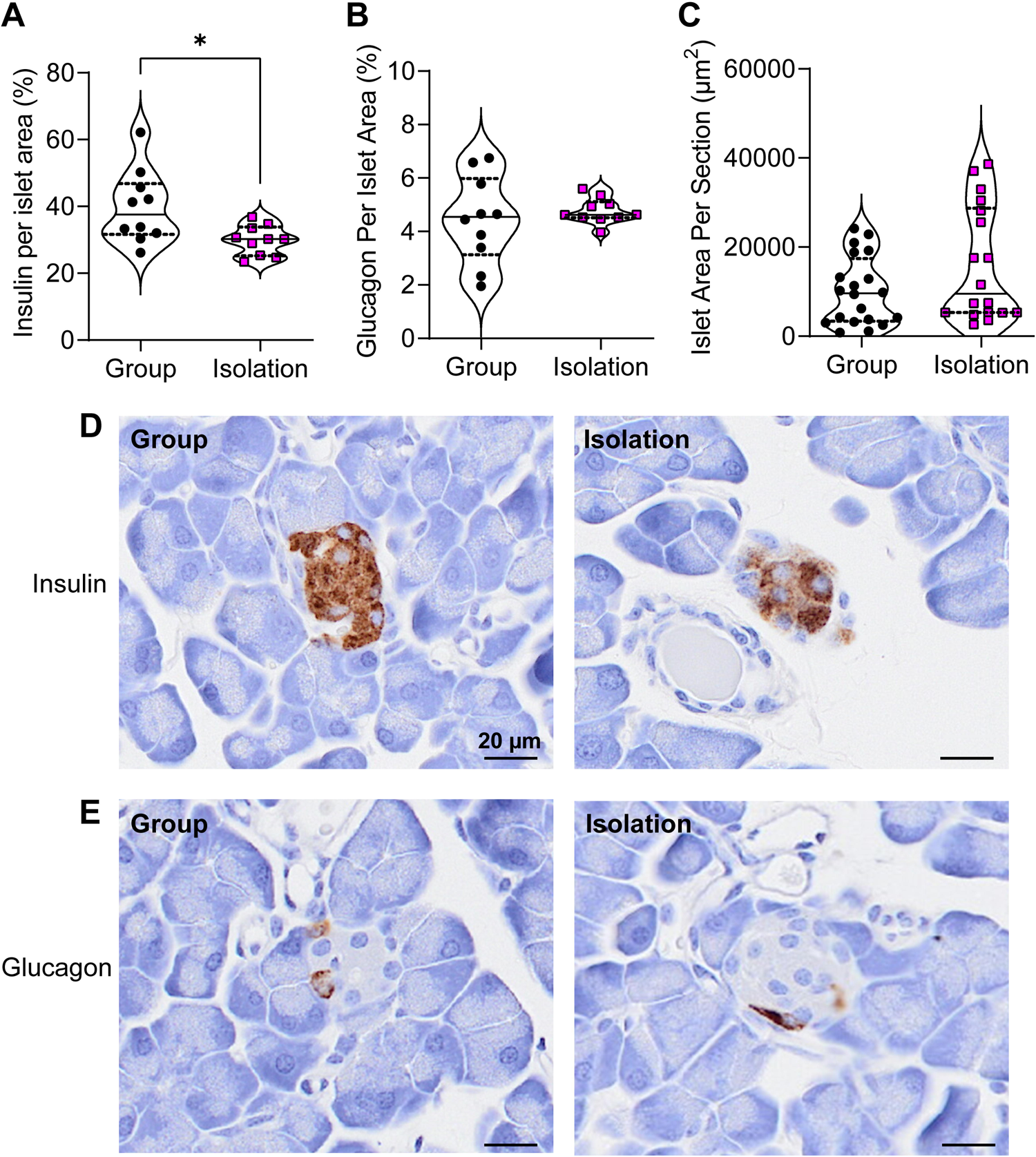
Comparison of pancreatic staining for insulin and glucagon. ***A:*** Percentage of total islet area within a pancreatic section that was found to be immunopositive for insulin. ***B:*** Same as ***A***, but for glucagon. ***C:*** Comparison of total islet area per pancreatic section. ***D:*** Representative insulin-stained islet for group-housed mouse (***left***) and adolescent-isolation mouse (***right***). ***E:*** Same as ***D***, but for glucagon. \**p* < 0.05

### Adolescent isolation induces a deficit in social behavior and negatively affects pain threshold

Since our adolescent-isolation paradigm directly removes social interaction with cage mates, we investigated any change in social interaction behaviors. Using a standard 3-chamber apparatus, we found the adolescent-isolation mice spent less total time interacting with a stranger mouse compared with group-housed controls (*p* = 0.0024) (**Figure 4A**). We did not observe a difference in the number of interactions with the enclosure, or the total number of entries to either chamber (**Figure 4B-C**). Adolescent isolation has been shown to have no effect on thermal sensitivity in ddY and CFW mouse strains, while increasing mechanical pain thresholds (41,42). In C57 mice, we also observed no difference in thermal sensitivity by measuring paw withdrawal time during a Hargreaves test (**Figure 4D**). Interestingly, we observed a decrease in mechanical pain thresholds in isolated C57 mice, opposite to that shown in ddY and CFW strains. Using a von Frey assessment, we found that isolated mice may have a reduction in 50% withdrawal threshold (*p* = 0.0503) and shallower average slope across filaments (*p* = 0.0235) (**Figure 4E-F**). Further 2-way analysis revealed a Condition x Filament interaction (*p* = 0.0193) (**Figure 4G**).

**Figure 4.**
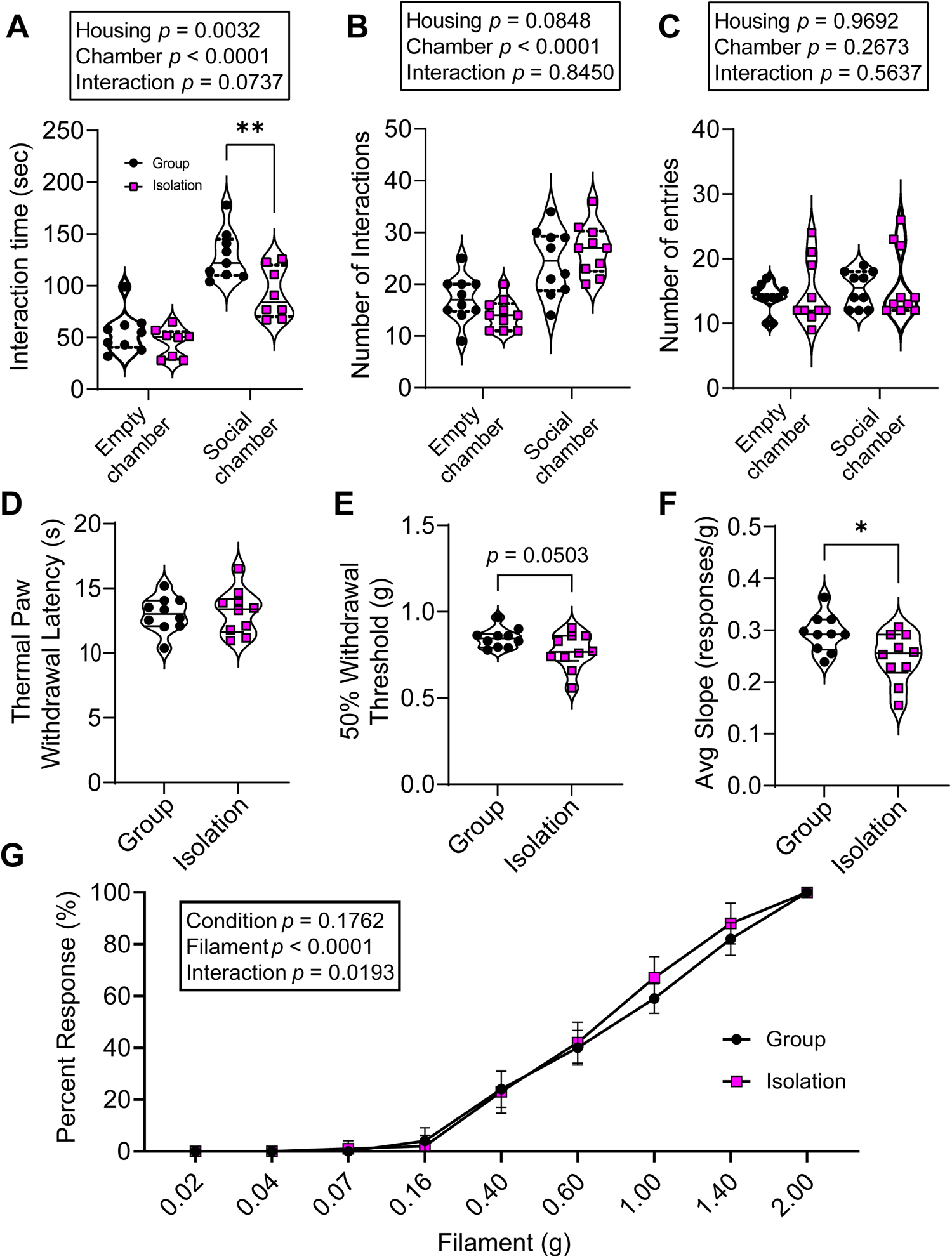
Social interaction (***A-C***) and nociceptive behaviors (***D-G***) ***A:*** Total interaction time with metal enclosure that contained no mouse (empty) or stranger mouse (social). ***B:*** Total number of interactions with metal enclosure that contained either no mouse or stranger mouse. ***C:*** Total number of entries to either the empty chamber or the social chamber (containing stranger mouse). ***D:*** Hargreaves test comparing thermal paw withdrawal latency. ***E:*** Von Frey test comparison of 50% withdrawal threshold. ***F:*** Average slope of responsiveness for von Frey test. ***G:*** Same as ***F***, but using a two-way comparison with plungers representing SEM. \**p* < 0.05, \*\**p* < 0.01

### Adolescent isolation affects food intake and sleep

To determine eating, drinking, and sleeping behaviors, mice were moved to Promethion metabolic phenotyping chambers. These chambers require mice to be individually housed, so this cohort was not used for further experiments after Promethion data collection. During the dark phase, adolescent-isolation mice consumed more chow (*p* = 0.0027) (**Figure 5A**) and spent less time asleep than mice that were previously group housed (*p* = 0.0044) (**Figure 5D**). There was no difference between groups in total water intake or locomotion during the dark phase (**Figure 5B-C**). Interestingly, we did observe a Condition x Time interaction for RER across the dark phase (*p* = 0.0056) (**Figure 5E**). During the light phase, we observed no difference in food intake, total water intake, locomotion, or RER (**Figure 5F-H, J**). However, adolescent-isolation mice spent less time asleep than mice that were previously group housed (*p* = 0.0006) (**Figure 5I**). We further found no differences between groups during the dark or light phase for oxygen consumption or thermogenesis, but adolescent-isolation mice spent proportionally less time resting inside their habitat than controls (**Supplemental Figure 1**).

**Figure 5.**
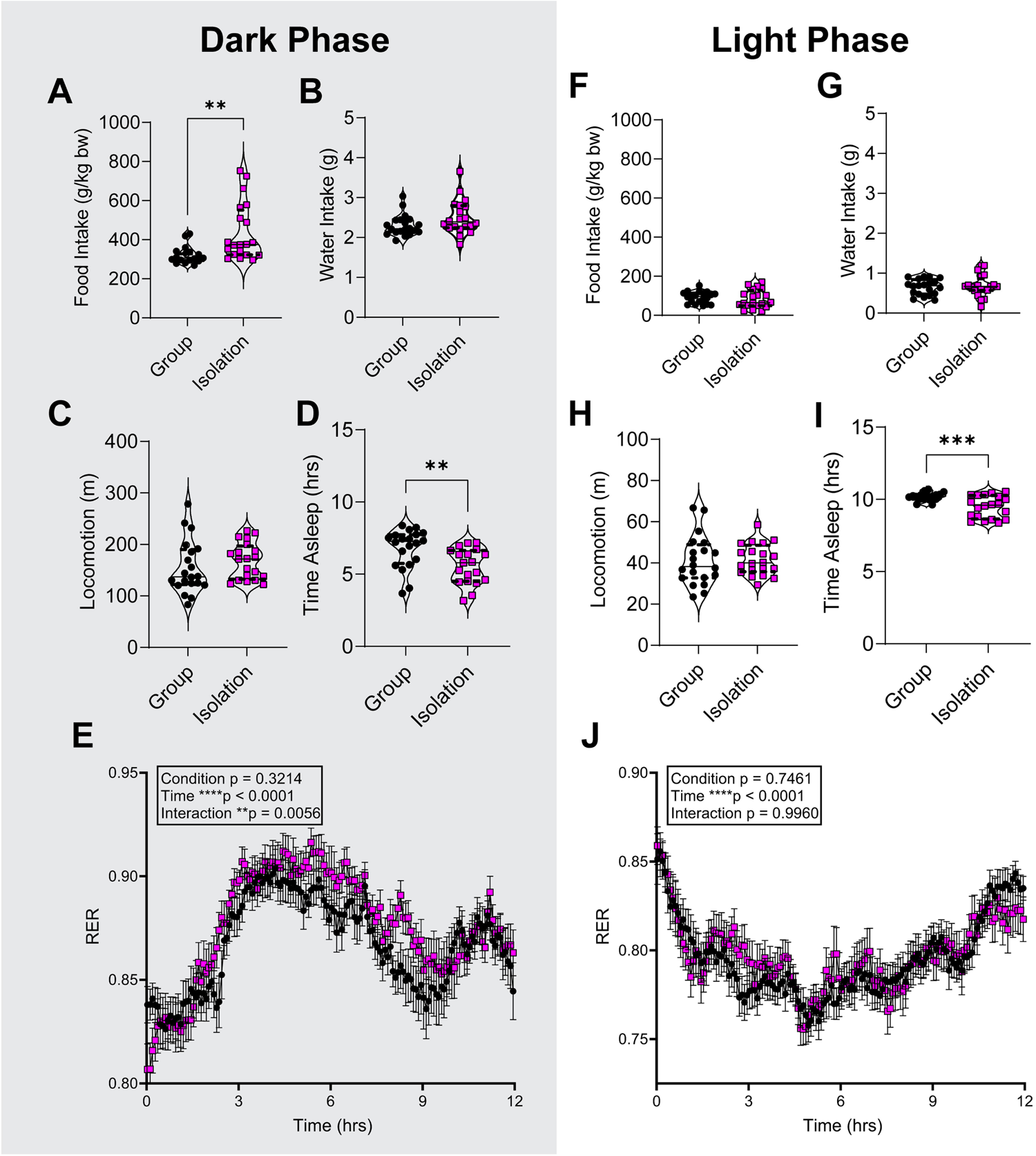
Assessment using Promethion metabolic phenotyping chambers separated by dark phase (***A-E***) and light phase (***F-J***). Each data point represents one light/dark cycle per mouse. ***A:*** Comparison of food intake relative to bodyweight for mice that were previously group-housed (Group) or isolated throughout adolescence (Isolation). ***B:*** Total water intake per dark phase. ***C:*** Total locomotion. ***D:*** Total time spent asleep throughout the dark phase, as defined by Promethion software. ***E:*** Respiratory exchange ratio (RER) throughout the dark phase. ***F-J:*** Same as ***A-E***, but for data collected during the light phase. \*\**p* < 0.01 \*\*\**p* < 0.001

### Isolation-induced type 2 diabetes is associated with changes in the neural 5-HT, GLP-1, and endocannabinoid circuits

To probe changes in neural gene expression that could be accompanying the phenotypic changes observed in adolescent-isolation mice, we used RT-qPCR to examine the olfactory bulb (OB), laterodorsal tegmental nucleus (LDT), lateral hypothalamic area (LHA), suprachiasmatic nucleus (SCN), arcuate nucleus and ventromedial hypothalamus (ARC/VMH), periaqueductal grey (PAG), and rostral ventromedial medulla (RVM). These regions were selected for their known roles in eating behavior, regulation of circadian behaviors, and nociception. We observed broad transcriptional changes in 5-HT-related genes (*Slc6a4*, *Htr1a*, *Htr2c*), insulinotropic genes (*Gcg*, *Insr*, *Glp1r*), and endocannabinoid genes (*Faah*, *Cnr1*) (**Figure 6**). The *Gcg* gene generates tissue-specific products, first encoding the PPG protein. In the pancreas, PPG is cleaved to create glucagon. In the neuropod cells of the gut and in PPGNs of the brain, PPG is cleaved and amidated to generate glucagon-like peptide 1 (GLP-1).

**Figure 6.**
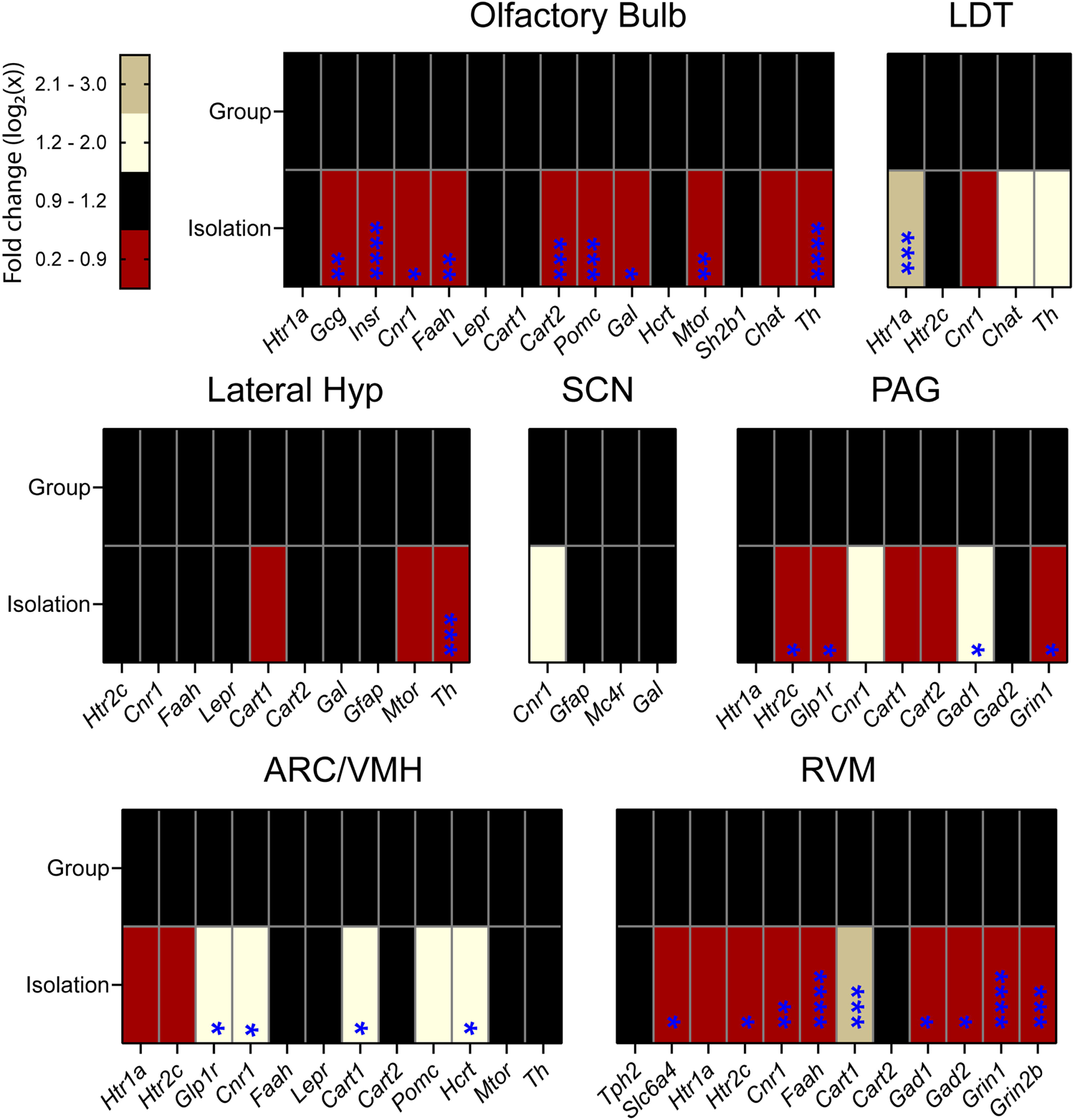
RT-qPCR of target genes. Relative changes in transcription are in adolescent-isolation mice (Isolation) comparison to group-housed mice (Group) and are represented using the color-coded heat map for fold change (left). Asterisks represent statistical significance of change, independent of effect size. Brain regions included the olfactory bulb, laterodorsal tegmental nucleus (LDT), lateral hypothalamic area (Lateral Hyp), suprachiasmatic nucleus (SCN), periaqueductal grey (PAG), arcuate nucleus with ventromedial hypothalamus (ARC/VMH), and rostral ventromedial medulla (RVM). \**p* < 0.05, \*\**p* < 0.01, \*\*\**p* < 0.001, \*\*\*\**p* < 0.0001

Since we observed changes in several genes associated with 5-HT and GLP-1 across many brain areas, we next used RNAscope (a multiplex *in situ* hybridization system) to spatially determine differences in the transcription of genes associated with the 5-HT/GLP-1 circuit. In the raphe nuclei (producing 5-HT), these included *Tph2* (which regulates 5-HT production) and *Glp1r* (GLP-1R). Activation of the GLP-1R on 5-HTNs is known to increase their activity, modulating appetite (25). We first profiled each raphe nucleus separately for the number of *in situ Glp1r* probe spots per *Tph2*-positive cell, then analyzed all raphe nuclei together. We found that the B9 5-HTNs of adolescent-isolation mice transcribed fewer copies of *Glp1r* compared to group-housed mice (*p* = 0.0227), and this held true for 5-HTNs across all raphe nuclei pooled together (*p* = 0.0447) (**Figure 7D-E**). We found no change in the relative percentage of *Tph2*-positive cells that expressed *Glp1r*, within each or across all raphe nuclei, respectively (**Figure 7F-G**).

**Figure 7.**
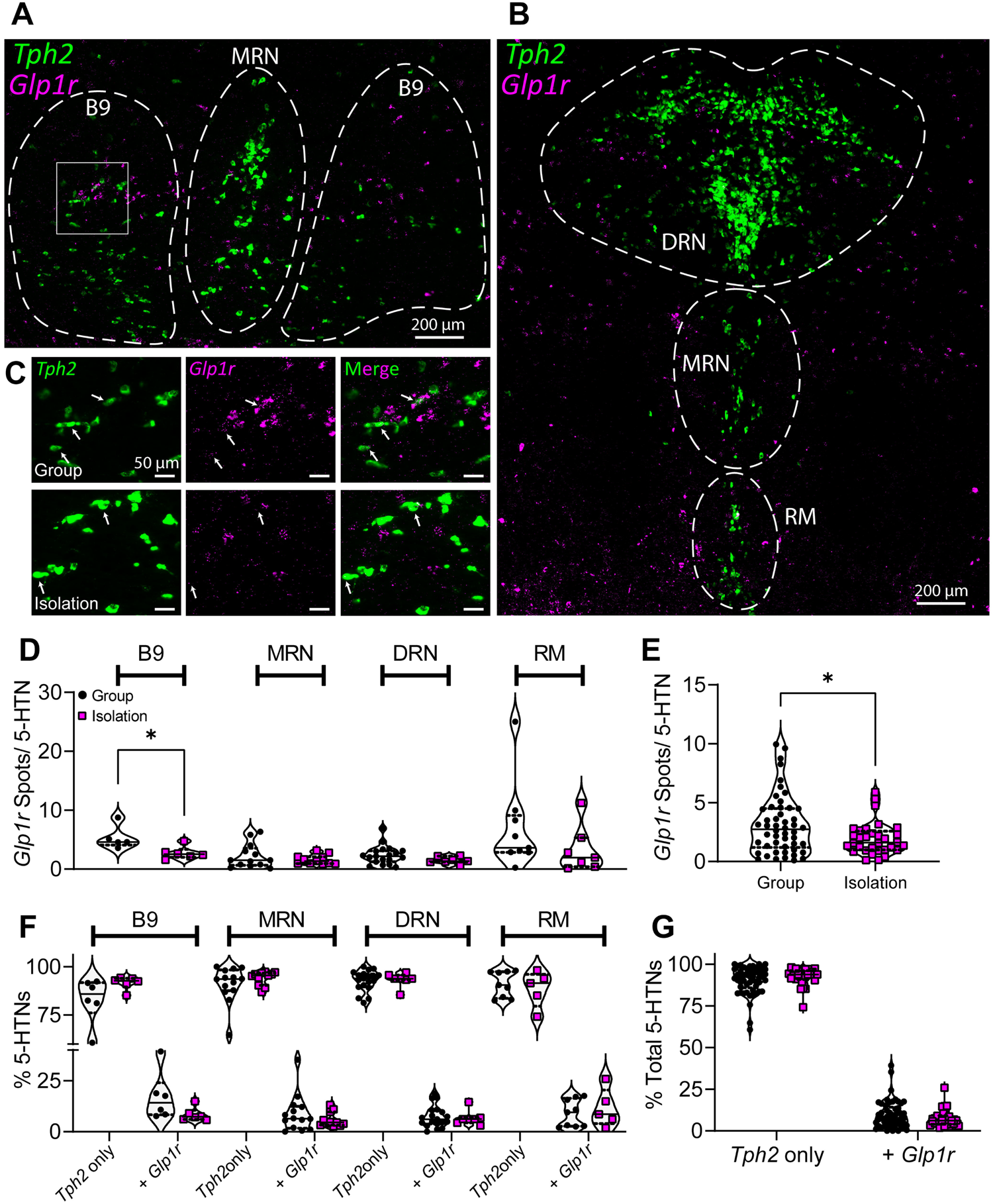
RNAscope assessment of the serotonergic nuclei. ***A:*** Coronal section of the rostral median raphe nucleus (MRN) and B9 serotonergic neurons, labeled with *in situ* probes for *Tph2* (green) and *Glp1r* (magenta). ***B:*** Same as ***A***, but including the dorsal raphe nucleus (DRN) and raphe magnus (RM). ***C:*** Channel-separated representative images for group-housed (***top***) and isolated (***bottom***) mice. ***Top*** row is taken from square region of interest in ***A***. ***D:*** Quantitation of *in situ* probe spots per *Tph2*-positive cell, group (black) vs isolation (magenta) separated by serotonergic nucleus. Each data point represents the mean across an entire tissue section. ***E:*** Same as ***D***, but combined across all anatomical regions. ***F:*** Percentage of *Tph2-*positive cells within each serotonergic nucleus with or without co-expression of *Glp1r*. ***G:*** Same as ***F***, but combined across all anatomical regions. \**p* < 0.05

To complement our findings in 5-HTNs, we spatially profiled transcription of *Gcg* (GLP-1), *Htr1a* (5-HT_1A_), and *Htr2c* (5-HT_2C_) in the PPGNs (those producing GLP-1). Activation of 5- HT_1A_ is known to inhibit PPGNs, and activation of 5-HT_2C_ is known to excite them (24). *Gcg*- positive cells of adolescent-isolation mice trended toward increased *Gcg* transcription (*p* = 0.0624), while exhibiting a decrease in *Htr1a* (*p* = 0.0227), and no change in *Htr2c* (**Figure 8C**). We found no change in the relative percentage of *Gcg*-positive cells that expressed either *Htr1a*, *Htr2c*, or both (**Figure 8D**).

**Figure 8.**
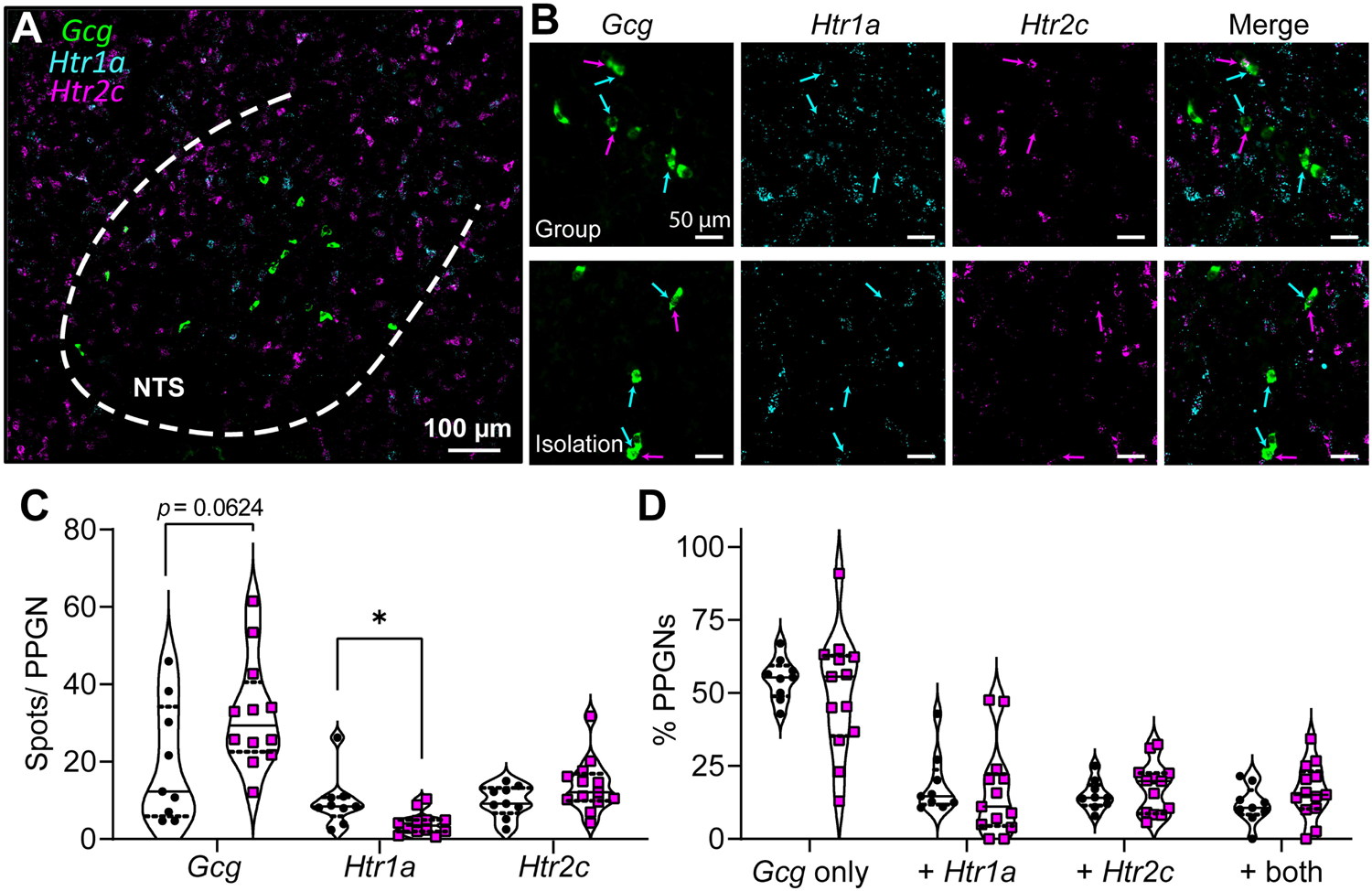
RNAscope assessment of PPG neurons of the brain stem. ***A:*** Coronal section nucleus of the solitary tract (NTS), labeled with *in situ* probes for *Gcg* (green), *Htr1a* (cyan) and *Htr2c* (magenta). ***B:*** Channel-separated representative images for group-housed (***top***) and isolated (***bottom***) mice. ***C:*** Quantitation of *in situ* probe spots per *Gcg*-positive cell, group (black) vs isolation (magenta). ***D:*** Percentage of *Gcg-*positive cells with or without co-expression of *Htr1a*, *Htr2c*, or both. \**p* < 0.05

## Discussion

This study reports that isolating C57 mice throughout adolescence is sufficient to induce type 2 diabetes per current definitions (39) and suggests the involvement of a neural 5-HT/GLP-1 circuit in this process. We report changes in 5-HT-related and insulinotropic genes in several brain regions, as well as complimentary changes in 5-HT and GLP-1 receptor genes within their respective nuclei. Our finding of type 2 diabetes was further associated with diminished insulin signaling, decreased insulin production, increased nociception, dysregulated sleep and eating behavior, and diminished plasma corticosterone.

It was previously found that isolating adolescent mice for 2 weeks caused a stark increase in food intake, while adult mice were unaffected by this paradigm (43). Additionally, isolating C57 mice from ages P35-P56 was not sufficient to induce obesity or diabetes (44). Our findings suggest that isolating mice throughout the entirety of adolescence may be necessary to induce a diabetic phenotype. Although our mice were only isolated for a period of 8 weeks, we found that pancreatic islets were less immunopositive for insulin. This suggests that type 2 diabetes may have progressed so far as to impair β-cell function (45). Interestingly, we observed diminished plasma corticosterone in the adolescent-isolation mice. This finding agrees with that found in long-term isolation models, wherein dysregulation of glucocorticoids may present as diminished basal levels that are hyper-elevated in response to stressors (46). Interestingly, our open-field assay did not suggest any anxiety-like behaviors in the adolescent-isolation mice (**Supplemental Figure 2**).

Peripheral 5-HT has long been implicated as a therapeutic target for dysregulated eating behaviors, but less is known about the role of neural 5-HT in type 2 diabetes and how it may be associated with insulinotropic hormones. Production of neural 5-HT is under the control of a different regulatory gene than peripheral 5-HT, and 5-HT cannot readily cross the blood-brain barrier. GLP-1 has recently gained popularity as a therapeutic to combat type 2 diabetes (47–49), which is produced centrally by the PPGNs of the NTS (and to a lesser extent the OB). 5-HT has been shown to differentially modulate PPGNs via 5-HT_1A_ and 5-HT_2C_ receptors, respectively (24). PPGNs of the NTS reciprocally project to the raphe nuclei, reducing appetite via GLP-1Rs that increase the activity of 5-HTNs (25). In adolescent-isolation mice, we find that 5-HTNs of the raphe nuclei produce fewer transcripts for the GLP-1R, and these mice consume more chow than group-housed controls. In parallel, PPGNs of the NTS produced fewer transcripts for the 5- HT_1A_ receptor, whose activation is known to inhibit the activity of these neurons (24). Taken together, these findings suggest an overall trend toward decreased 5-HTN activity (via reduction of GLP-1-mediated excitation) and increased PPGN activity (via decreased 5-HT-mediated inhibition). This is supported by a previous report that the activity of 5-HTNs was strongly reduced in response to electrophysiological, optical, and neuromodulatory stimuli following an adolescent-isolation paradigm (23). This 5-HT/GLP-1 circuit may represent an intersectional target to further investigate the relationship between social isolation and type 2 diabetes, as well as a pharmacologically-relevant circuit to explore the effects of serotonin and GLP-1 receptor agonists.

Further investigation is needed to determine if female mice undergo the same neural changes, even though they don’t develop peripheral diabetes in the same timeline (29). This could determine whether isolation-induced changes in neural gene expression are caused by the peripheral changes in insulin sensitivity, or if these neural changes precede peripheral type 2 diabetes.

## Acknowledgements

All authors critically evaluated the work for important intellectual content. All authors acquired, analyzed, or interpreted data in the study. As such, we confirm that all authors approved the final version of the manuscript, agreed to be accountable for all aspects of the work, that all persons designated as authors qualify for authorship, and all those who qualify for authorship are listed. In using the CRediT (Contributor Roles Taxonomy) guidelines, L.K., K.K., N.B. and C.M. were responsible for project conceptualization, formal analyses, and methodology. L.K., K.K., N.B. and D.G. performed data acquisition and analysis. L.K. was responsible for writing the original draft of the manuscript, all authors contributed to reviewing and editing the manuscript. C.M. and K.R. were responsible for project administration and supervision. C.M. is the guarantor of this work and, as such, has had full access to all data and takes responsibility for its integrity and analysis.

We thank Yu Xu for performing routine technical assistance and maintaining mouse colonies, Eric Weatherford of the Metabolic Phenotyping Core Laboratory for operating the Promethion system, Samantha Pierson for contributing to figure design, as well as Mariah Leidinger of the Comparative Pathology Laboratory for performing immunostaining pancreata.

