## Supplementary Material for "Involvement of a serotonin/GLP-1 circuit in adolescent isolation-induced diabetes"

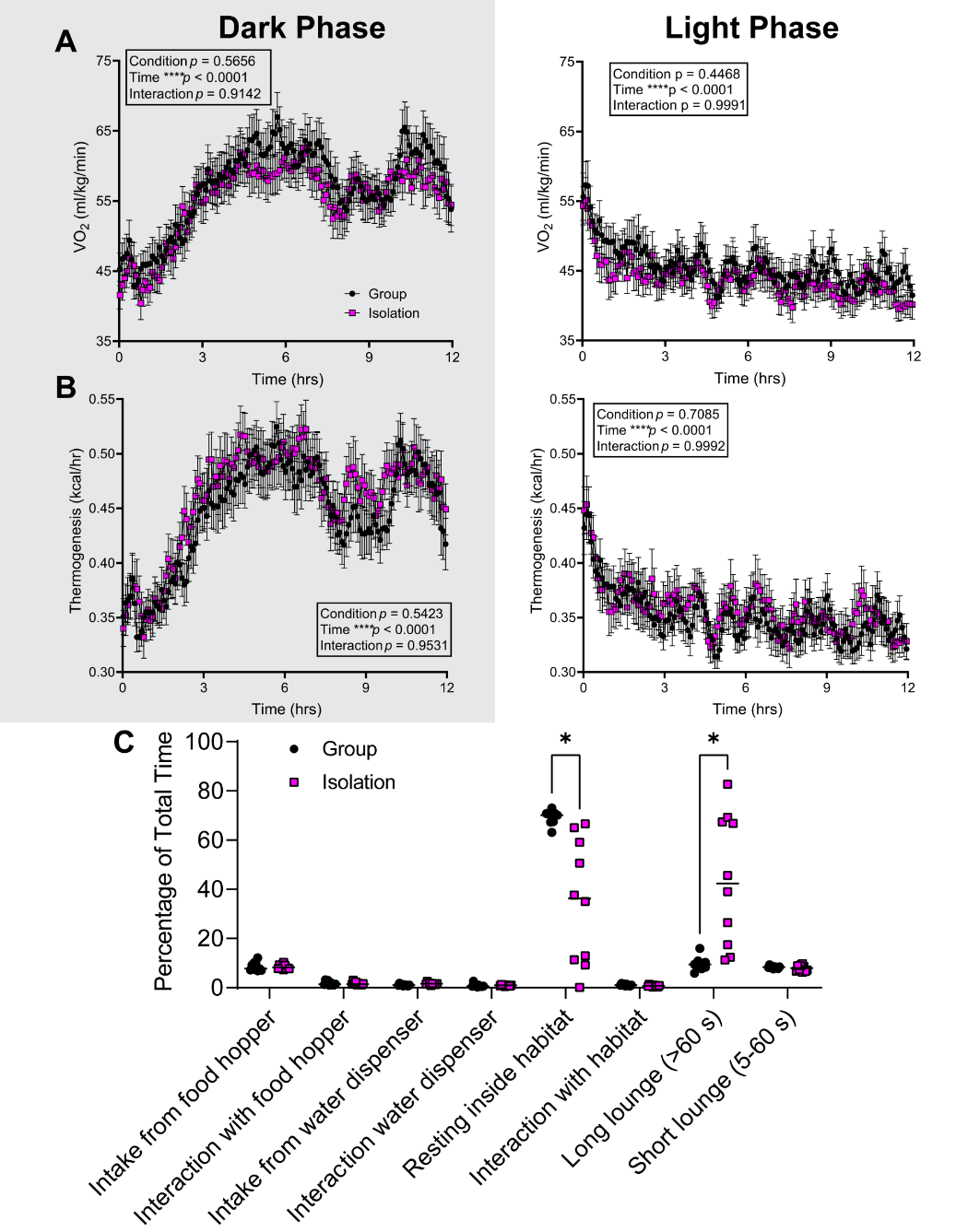
**SUPPLEMENTARY FIGURES:**

**Supplemental Figure 1** Metabolic assessment using Promethion. ***A:*** Relative oxygen consumption (VO2) during the dark phase (left) and light phase (right). ***B:*** Thermogenesis during the dark phase (left) and light phase (right). ***C:*** Cumulative time budget for entire measurement period. “Resting inside habitat” and “long lounge” are periods of non-movement which persist for >60 s.

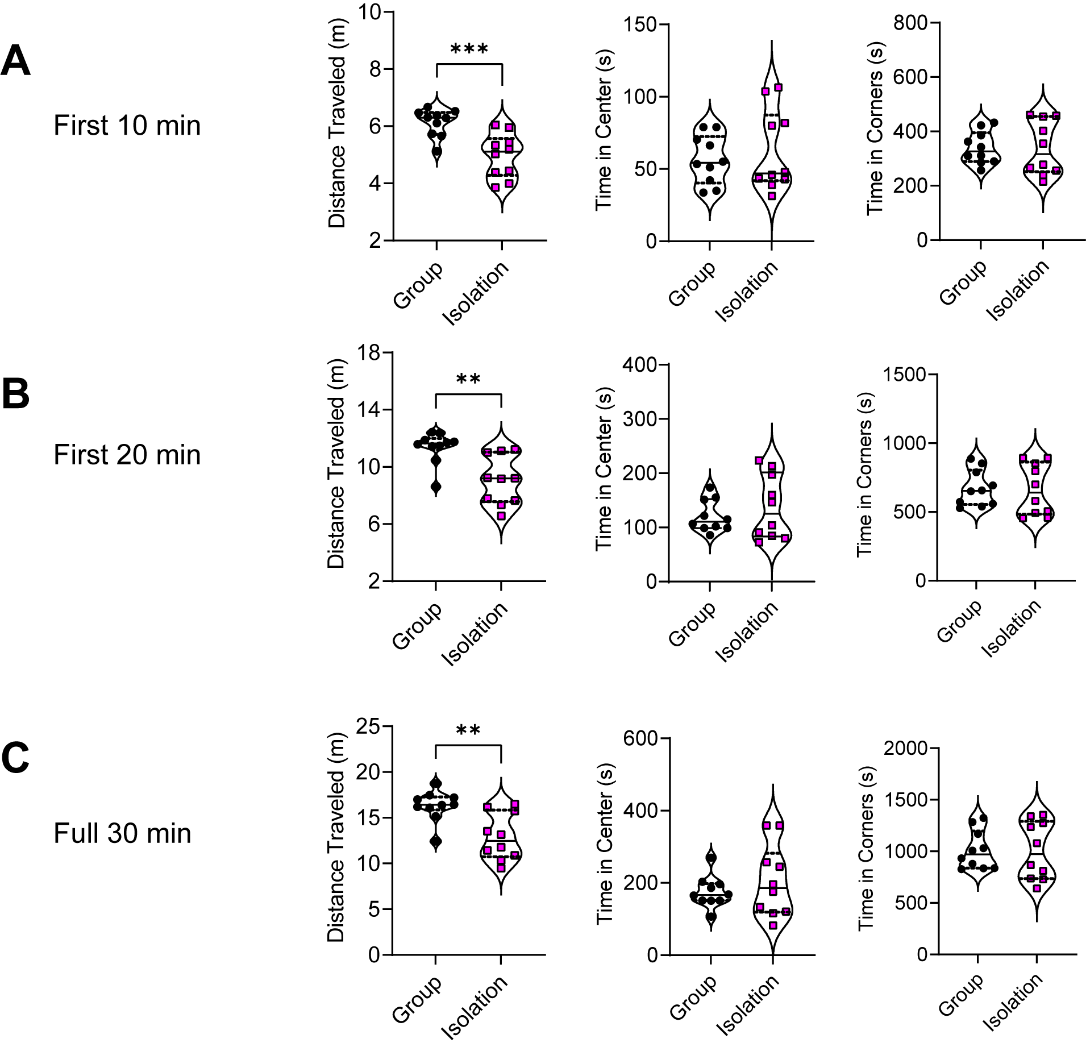

**Supplemental Figure 2 *A:*** Total distance traveled during open-field exploration task (***left***), total time spent in center of arena (***center***) and total time spent in corner areas (***right***) during first 10 minutes of test. ***B:*** Same as (***A***), but first 20 minutes of test. ***C:*** Same as (***A***), but full 30 minutes of test.

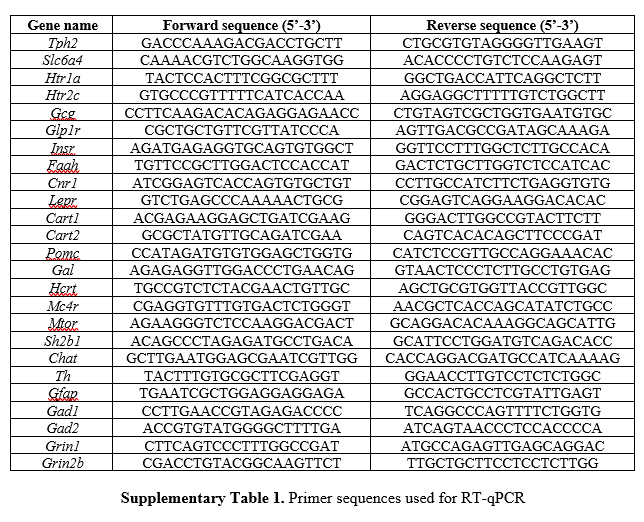

**SUPPLEMENTARY METHODS:**

Open field test: Mice were placed in the corner of a 50 x 50 x 25 cm plexiglass arena (20 lux) and allowed to freely explore the arena for 30 min. The open field test was performed to evaluate locomotor and exploratory behavior. The total distance traveled (cm), time spent in the center of the arena, and time spent in the corners of the arena were recorded by an overhead camera and quantified using Ethovision XT14. The center of the open field was defined as the central 15% of the arena.

**FULL STATISTICS:**

| **Figure No.** | | | **Experiment** | | | | **Mean ± SD (n)** | | | | **Statistical test, t, df, *p*** |
| --- | --- | --- | --- | --- | --- | --- | --- | --- | --- | --- | --- |
| Figure 1B | | | Blood glucose comparison | | | | Group = 117.8 ± 21.7 mg/dl (10) vs Isolation = 208.5 ± 34.28 mg/dl (10) | | | | Student’s *t*-test, 7.063, 18, *****p* < 0.0001 |
| Figure 1C | | | Glucose tolerance test | | | | Group = 34265 ± 4712 (10) vs Isolation = 37308 ± 5368 (10) | | | | Student’s *t*-test, 1.347, 18, *p* = 0.1946 |
| Figure 1D | | | Insulin tolerance test | | | | Group = 7923 ± 2340 (9) vs Isolation = 15763 ± 3521 (10) | | | | Student’s *t*-test, 5.644, 17, *****p* < 0.0001 |
| Figure 1E | | | Plasma insulin | | | | Group = 17.13 ± 12.31 mIU/ml (9) vs Isolation = 41.00 ± 46.94 mIU/ml (10) | | | | *t*-test with Welch’s correction, 1.550, 10.36, *p* = 0.1511 |
| Figure 1F | | | Plasma leptin | | | | Group = 2.007 ± 0.981 pg/ml (9) vs Isolation = 2.679 ± 1.579 pg/ml (10) | | | | Student’s *t*-test, 1.097, 17, *p* = 0.2878 |
| Figure 1G | | | Plasma corticosterone | | | | Group = 4699 ± 2489 pg/ml (10) vs Isolation = 2281 ± 3299 pg/ml (10) | | | | Student’s *t*-test, 4.236, 16, ****p* = 0.0006 |
| Figure 1H | | | Body weight comparison | | | | Group = 26.91 ± 2.49 g (10) vs Isolation = 25.74 ± 1.10 g (10) | | | | Mann-Whitney U, *p* = 0.6031 |
| Figure 1I | | | % lean tissue | | | | Group = 68.22 ± 3.09 % (10) vs Isolation = 65.37 ± 1.94 % (10) | | | | Student’s *t*-test, 2.475, 18, **p* = 0.0235 |
| Figure 1J | | | % free body fluid | | | | Group = 5.811 ± 0.472 % (10) vs Isolation = 5.264 ± 0.316 % (10) | | | | Student’s *t*-test, 3.045, 18, ***p* = 0.0070 |
| Figure 1K | | | % body fat | | | | Group = 10.19 ± 3.73 % (10) vs Isolation = 12.79 ± 2.53 % (10) | | | | Student’s *t*-test, 1.830, 18, *p* = 0.0839 |
| **Figure No.** | | | | **Experiment** | | | | **Statistical Test, F stat** | | | **Multiple comparisons** |
| Figure 2A | | | | liver | | | | ANOVA, F (3, 15) = 2.069 | | | Sidak’s multiple comparison test (Group insulin vs Isolation insulin) *p* = 0.6505 |
| Figure 2B | | | | muscle | | | | ANOVA, F (3, 15) = 2.262 | | | Sidak’s multiple comparison test (Group insulin vs Isolation insulin) *****p* < 0.0001 |
| Figure 2C | | | | BAT | | | | ANOVA, F (3, 15) = 0.6086, | | | Sidak’s multiple comparison test (Group insulin vs Isolation insulin) *p* = 0.2814 |
| Figure 2D | | | | WAT | | | | ANOVA, F (3, 15) = 2.206 | | | Sidak’s multiple comparison test (Group insulin vs Isolation insulin) *p* = 0.7834 |
| **Figure No.** | | | | **Experiment** | | | | **Mean ± SD (n)** | | | **Statistical test, t, df, *p*** |
| Figure 3A | | | | % islet area immunopositive for insulin | | | | Group = 39.72 ± 10.87 % (10) vs Isolation = 29.87 ± 4.421 % (10) | | | *t*-test with arcsine transformation, 2.671, 18, **p* = 0.0156 |
| Figure 3B | | | | % glucagon area immunopositive for insulin | | | | Group = 4.439 ± 1.627 % (10) vs Isolation = 4.765 ± 0.473 % (10) | | | *t*-test with arcsine transformation, 0.8136, 10.30, *p* = 0.4343 |
| Figure 3C | | | | Total islet area per section | | | | Group = 10113 ± 7622 µm^2^ (10) vs Isolation = 15931 ± 9572 µm^2^ (10) | | | Mann-Whitney U, *p* = 0.1557 |
| **Figure No.** | **Experiment** | | | | | | | **Statistical Test, F stat** | | | **Multiple comparisons** |
| Figure 4A | Interaction time social/empty enclosure | | | | | | | 2-w mixed RM ANOVA, Housing Condition F (1, 30) = 10.25, ***p* = 0.0032; Chamber F (1, 30) = 70.22, *****p* < 0.0001; Housing x Chamber interaction F (1, 30) = 3.435, *p* = 0.0737, | | | Sidak’s multiple comparison test (Group Social chamber vs Isolation Social chamber) ***p* = 0.0024 |
| Figure 4B | Number of interactions with social/empty enclosure | | | | | | | 2-w mixed RM ANOVA, Housing Condition F (1, 36) = 0.0388, *p* = 0.8450; Chamber F (1, 36) = 41.39, *****p* < 0.0001; Housing x Chamber interaction F (1, 36) = 3.141, *p* = 0.0848 | | |  |
| Figure 4C | Number of entries to social/empty chamber | | | | | | | 2-w mixed RM ANOVA, Housing Condition F (1, 36) = 0.3396, *p* = 0.5637; Chamber F (1, 36) = 1.269, *p* = 0.2673; Housing x Chamber interaction F (1, 36) = 0.0015, *p* = 0.9692 | | |  |
| **Figure No.** | **Experiment** | | | | | | | **Mean ± SD (n)** | | | **Statistical test, t, df, *p*** |
| Figure 4D | Hargreaves | | | | | | | Group = 12.99 ± 1.35 s (10) vs Isolation = 13.19 ± 1.72 s (10) | | | Student’s *t*-test, 0.2790, 18, *p* = 0.7834 |
| Figure 4E | 50% withdraw threshold | | | | | | | Group = 0.8474 ± 0.0572 g (10) vs Isolation = 0.7684 ± 0.1045 g (10) | | | Student’s *t*-test, 2.098, 18, *p* = 0.0503 |
| Figure 4F | Average response slope | | | | | | | Group = 0.2940 ± 0.0362 responses/g (10) vs Isolation = 0.2469 ± 0.0481 responses/g (10) | | | Student’s *t*-test, 2.474, 18, **p* = 0.0235 |
| **Figure No.** | **Experiment** | | | | | | | **Statistical Test, F stat** | | | **Multiple comparisons** |
| Figure 4G | Percent response v Filament | | | | | | | 2-w mixed RM ANOVA, Housing Condition F (1, 18) = 1.982, *p* = 0.1762; Filament F (3.713, 66.83) = 1334, *****p* < 0.001; Condition x Filament interaction F (8, 144) = 2.384, **p* = 0.0193 | | |  |
| **Figure No.** | | **Experiment** | | | | | **Mean ± SD (n)** | | | | **Statistical test, t, df, *p*** |
| Figure 5A | | Food consumption dark phase | | | | | Group = 321.4 ± 45.6 g/kg (17) vs Isolation = 437.3 ± 146.1 g/kg (20 | | | | *t*-test with Welch’s correction, 3.361, 23.24, ***p* = 0.0027 |
| Figure 5B | | Water intake dark phase | | | | | Group = 2.30 ± 0.27 g (20) vs Isolation = 2.51 ± 0.42 g (20) | | | | Student’s *t*-test, 1.822, 38, *p* = 0.0763 |
| Figure 5C | | Locomotion dark phase | | | | | Group = 155.0 ± 51.7 m (20) vs Isolation = 168.1 ± 35.1 m (20) | | | | Student’s *t*-test, 0.9343, 38, *p* = 0.3561 |
| Figure 5D | | Time asleep dark phase | | | | | Group = 6.80 ± 1.34 h (20) vs Isolation = 5.55 ± 1.22 h (19) | | | | Student’s *t*-test, 3.034, 37, ***p* = 0.0044 |
| **Figure No.** | | **Experiment** | | | | | **Statistical Test, F stat** | | | | **Multiple comparisons** |
| Figure 5E | | RER dark phase | | | | | 2-w mixed RM ANOVA, Dark: Condition x Time F (143, 5434) = 1.331, ***p* = 0.0056 | | | |  |
| **Figure No.** | | **Experiment** | | | | | **Mean ± SD (n)** | | | | **Statistical test, t, df, *p*** |
| Figure 5F | | Food consumption light phase | | | | | Group = 92.2 ± 29.5 g/kg (20) vs Isolation = 84.1 ± 46.8 g/kg (20) | | | | Student’s *t*-test, 0.6538, 38, *p* = 0.5172 |
| Figure 5G | | Water intake light phase | | | | | (Group = 0.65 ± 0.20 g (20) vs Isolation = 0.68 ± 0.27 g (20) | | | | Student’s *t*-test, 0.4587, 38, *p* = 0.6491 |
| Figure 5H | | Locomotion light phase | | | | | Group = 41.0 ± 12.1 m (20) vs Isolation = 41.6 ± 7.6 m (20) | | | | Student’s *t*-test, 0.1638, 38, *p* = 0.8708 |
| Figure 5I | | Time asleep light phase | | | | | Group = 10.19 ± 0.28 h (18) vs Isolation = 9.48 ± 0.76 h (20) | | | | *t*-test with Welch’s correction, 3.918, 24.49, ****p* = 0.0006 |
| **Figure No.** | | **Experiment** | | | | | **Statistical Test, F stat** | | | | **Multiple comparisons** |
| Figure 5J | | RER light phase | | | | | 2-w mixed RM ANOVA, Light: Housing Condition F (1, 38) = 1.010, *p* = 0.3214; Light: Housing Condition F (1, 38) = 0.1064, *p* = 0.7461) | | | |  |
| **Figure No.** | | | | | **Brain region** | **Primer** | | | **Fold change** (log_2_(x)) | **t ratio, df, *p*** (Multiple unpaired t-tests, Benjamini, Krieger, and Yekutieli with assumption of individual variance for each row) | |
| Figure 6 | | | | | **Olfactory bulb** | *Htr1a* | | | -0.0745 | 0.8733, 11, *p* = 0.4011 | |
|  |  |  |  |  |  | *Gcg* | | | -0.3679 | 3.872, 13, ***p* = 0.0019 | |
|  |  |  |  |  |  | *Insr* | | | -0.4354 | 5.784, 12, *****p* < 0.0001 | |
|  |  |  |  |  |  | *Cnr1* | | | -0.2414 | 2.371, 13, **p* = 0.0339 | |
|  |  |  |  |  |  | *Faah* | | | -0.4041 | 3.952, 13, ***p* = 0.0017 | |
|  |  |  |  |  |  | *Lepr* | | | -0.0523 | 0.5694, 12, *p* = 0.5796 | |
|  |  |  |  |  |  | *Cart1* | | | -0.0827 | 0.8546, 11, *p* = 0.4110 | |
|  |  |  |  |  |  | *Cart2* | | | -0.4286 | 4.497, 12, ****p* = 0.0007 | |
|  |  |  |  |  |  | *Pomc* | | | -0.8458 | 5.726, 11, ****p* = 0.0001 | |
|  |  |  |  |  |  | *Gal* | | | -0.2404 | 2.871, 13, **p* = 0.0131 | |
|  |  |  |  |  |  | *Hcrt* | | | +0.0314 | 0.06907, 7, *p* = 0.9469 | |
|  |  |  |  |  |  | *Mtor* | | | -0.2387 | 4.151, 13, ***p* = 0.0011 | |
|  |  |  |  |  |  | *Sh2b1* | | | -0.0299 | 0.3775, 13, *p* = 0.7119 | |
|  |  |  |  |  |  | *Chat* | | | -0.2965 | 1.229, 11, *p* = 0.2446 | |
|  |  |  |  |  |  | *Th* | | | -0.5756 | 7.588, 13, *****p* < 0.0001 | |
|  |  |  |  |  | **LDT** | *Htr1a* | | | +1.0983 | 5.110, 10, ****p* = 0.0005 | |
|  |  |  |  |  |  | *Htr2c* | | | +0.1532 | 0.4353, 6, *p* = 0.6786 | |
|  |  |  |  |  |  | *Cnr1* | | | -0.7725 | 1.625, 6, *p* = 0.1552 | |
|  |  |  |  |  |  | *Chat* | | | +0.3391 | 1.203, 10, *p* = 0.2567 | |
|  |  |  |  |  |  | *Th* | | | +0.6544 | 1.043, 8, *p* = 0.3273 | |
|  |  |  |  |  | **Lateral Hyp** | *Htr2c* | | | +0.0328 | 0.4470, 12, *p* = 0.6628 | |
|  |  |  |  |  |  | *Cnr1* | | | -0.0289 | 0.6298, 13, *p* = 0.5398 | |
|  |  |  |  |  |  | *Faah* | | | +0.0718 | 0.2468, 12, *p* = 0.8092 | |
|  |  |  |  |  |  | *Lepr* | | | +0.2029 | 0.4271, 13, *p* = 0.6763 | |
|  |  |  |  |  |  | *Cart1* | | | -0.2455 | 1.855, 12, *p* = 0.0883 | |
|  |  |  |  |  |  | *Cart2* | | | -0.1470 | 1.261, 13, *p* = 0.2295 | |
|  |  |  |  |  |  | *Gal* | | | -0.0190 | 0.3900, 14, *p* = 0.7024 | |
|  |  |  |  |  |  | *Gfap* | | | -0.1502 | 1.139, 14, *p* = 0.2737 | |
|  |  |  |  |  |  | *Mtor* | | | -0.2939 | 1.467, 11, *p* = 0.1703 | |
|  |  |  |  |  |  | *Th* | | | -0.7075 | 5.021, 13, ****p* = 0.0002 | |
|  |  |  |  |  | **SCN** | *Cnr1* | | | +0.3885 | 1.856, 10, *p* = 0.0931 | |
|  |  |  |  |  |  | *Gfap* | | | +0.0510 | 0.1551, 10, *p* = 0.8798 | |
|  |  |  |  |  |  | *Mc4r* | | | +0.2216 | 1.053, 10, *p* = 0.3173 | |
|  |  |  |  |  |  | *Gal* | | | +0.1966 | 0.2685, 10, *p* = 0.7938 | |
|  |  |  |  |  | **PAG** | *Htr1a* | | | -0.0775 | 0.3648, 13, *p* = 0.7211 | |
|  |  |  |  |  |  | *Htr2c* | | | -0.3957 | 2.733, 13, **p* = 0.0171 | |
|  |  |  |  |  |  | *Glp1r* | | | -0.6936 | 2.792, 11, **p* = 0.0175 | |
|  |  |  |  |  |  | *Cnr1* | | | +0.5271 | 1.942, 13, *p* = 0.0741 | |
|  |  |  |  |  |  | *Cart1* | | | -0.2178 | 0.6651, 11, *p* = 0.5197 | |
|  |  |  |  |  |  | *Cart2* | | | -0.3722 | 1.002, 10, *p* = 0.3401 | |
|  |  |  |  |  |  | *Gad1* | | | +0.6772 | 2.608, 12, **p* = 0.0229 | |
|  |  |  |  |  |  | *Gad2* | | | +0.0510 | 0.002513, 13, *p* = 0.9980 | |
|  |  |  |  |  |  | *Grin1* | | | -0.2435 | 2.404, 14, **p* = 0.0306 | |
|  |  |  |  |  |  | *App* | | | -0.2740 | 2.735, 13, **p* = 0.0170 | |
|  |  |  |  |  | **ARC/VMH** | *Htr1a* | | | -0.2438 | 1.924, 10, *p* = 0.0833 | |
|  |  |  |  |  |  | *Htr2c* | | | -0.1830 | 1.151, 10, *p* = 0.2765 | |
|  |  |  |  |  |  | *Glp1r* | | | +0.5636 | 2.712, 9, **p* = 0.0239 | |
|  |  |  |  |  |  | *Cnr1* | | | +0.2940 | 2.993, 10, **p* = 0.0135 | |
|  |  |  |  |  |  | *Faah* | | | -0.0209 | 0.2717, 10, *p* = 0.7914 | |
|  |  |  |  |  |  | *Lepr* | | | +0.0854 | 0.2796, 10, *p* = 0.7855 | |
|  |  |  |  |  |  | *Cart1* | | | +0.3741 | 2.876, 10, **p* = 0.0165 | |
|  |  |  |  |  |  | *Cart2* | | | +0.1532 | 0.9508, 10, *p* = 0.3641 | |
|  |  |  |  |  |  | *Pomc* | | | +0.3138 | 0.6796, 10, *p* = 0.5122 | |
|  |  |  |  |  |  | *Hcrt* | | | +0.8718 | 3.085, 9, **p* = 0.0130 | |
|  |  |  |  |  |  | *Mtor* | | | +0.0058 | 0.008664, 10, *p* = 0.9933 | |
|  |  |  |  |  |  | *Th* | | | +0.0468 | 0.2443, 10, *p* = 0.8119 | |
|  |  |  |  |  | **RVM** | *Tph2* | | | -0.0988 | 0.8623, 12, *p* = 0.4054 | |
|  |  |  |  |  |  | *Slc6a4* | | | -0.5517 | 2.518, 11, **p* = 0.0286 | |
|  |  |  |  |  |  | *Htr1a* | | | -0.1696 | 0.8159, 12, *p* = 0.4304 | |
|  |  |  |  |  |  | *Htr2c* | | | -0.4317 | 2.669, 13, **p* = 0.0193 | |
|  |  |  |  |  |  | *Cnr1* | | | -0.3741 | 3.152, 13, ***p* = 0.0076 | |
|  |  |  |  |  |  | *Faah* | | | -0.5513 | 6.132, 13, *****p* < 0.0001 | |
|  |  |  |  |  |  | *Cart1* | | | +1.2394 | 4.171, 11, ****p* = 0.0016 | |
|  |  |  |  |  |  | *Cart2* | | | +0.0772 | 0.2596, 13, *p* = 0.7992 | |
|  |  |  |  |  |  | *Gad1* | | | -0.4272 | 2.511, 13, **p* = 0.0260 | |
|  |  |  |  |  |  | *Gad2* | | | -0.4822 | 2.623, 14, **p* = 0.0201 | |
|  |  |  |  |  |  | *Grin1* | | | -0.5494 | 7.110, 11, *****p* < 0.0001 | |
|  |  |  |  |  |  | *Grin2b* | | | -0.7063 | 4.468, 12, ****p* = 0.0008 | |

| **Figure No.** | **Experiment** | **Mean ± SD (n)** | **Statistical test, t, df, *p*** |
| --- | --- | --- | --- |
| Figure 7D | *Glp1r* spots/ 5HTN B9 | Group = 5.314 ± 1.992 (5) vs Isolation = 2.734 ± 1.081 (6) | Student’s *t*-test, 2.743, 9, **p* = 0.0227 |
|  | *Glp1r* spots/ 5HTN MRN | Group = 2.192 ± 2.071 (13) vs Isolation = 1.615 ± 0.811 (11) | Mann-Whitney U, *p* = 0.9095 |
|  | *Glp1r* spots/ 5HTN DRN | Group = 2.355 ± 1.579 (19) vs Isolation = 1.495 ± 0.595 (7) | Mann-Whitney U, *p* = 0.1516 |
|  | *Glp1r* spots/ 5HTN RM | Group = 6.839 ± 7.447 (9) vs Isolation = 3.289 ± 3.919 (7) | Mann-Whitney U, *p* = 0.1416 |
| Figure 7E | *Glp1r* spots/ 5HTN | Group = 3.122 ± 2.496 (50) vs Isolation = 1.457 ± 1.369 (32) | Mann-Whitney U, **p* = 0.0447 |
| Figure 7F | %5-HTNs B9 *Tph2* only | Group = 82.89 ± 11.82 % (6) vs Isolation = 91.61 ± 3.38 % (6) | Student’s *t*-test, 1.737, 10, *p* = 0.1130 |
|  | %5-HTNs B9 *Tph2* + *Glp1r* | Group = 17.11 ± 11.82 % (6) vs Isolation = 8.39 ± 3.38 % (6) | Student’s *t*-test, 1.737, 10, *p* = 0.1130 |
|  | %5-HTNs MRN *Tph2* only | Group = 93.21 ± 5.28 % (13) vs Isolation = 89.04 ± 2.85 % (10) | Student’s *t*-test, 0.8326, 21, *p* = 0.4144 |
|  | %5-HTNs MRN *Tph2* + *Glp1r* | Group = 6.79 ± 5.28 % (13) vs Isolation = 5.25 ± 2.85 % (10) | Student’s *t*-test, 0.8326, 21, *p* = 0.4144 |
|  | %5-HTNs DRN *Tph2* only | Group = 92.68 ± 5.33 % (19) vs Isolation = 93.23 ± 3.69 % (7) | Student’s *t*-test, 0.2524, 24, *p* = 0.8029 |
|  | %5-HTNs DRN *Tph2* + *Glp1r* | Group = 7.32 ± 5.33 % (19) vs Isolation = 6.77 ± 3.69 % (7) | Student’s *t*-test, 0.2524, 24, *p* = 0.8029 |
|  | %5-HTNs RM *Tph2* only | Group = 90.97 ± 6.62 % (9) vs Isolation = 88.52 ± 9.34 % (5) | Student’s *t*-test, 0.5749, 12, *p* = 0.5760 |
|  | %5-HTNs RM *Tph2* + *Glp1r* | Group = 9.03 ± 6.62 % (9) vs Isolation = 11.48 ± 9.34 % (5) | Student’s *t*-test, 0.5749, 12, *p* = 0.5760 |
| Figure 7G | %5-HTNs *Tph2* only | Group = 91.57 ± 6.17 % (52) vs Isolation = 93.19 ± 3.79 % (29) | Student’s *t*-test, 1.283, 79, *p* = 0.2032 |
|  | %5-HTNs *Tph2* + *Glp1r* | Group = 8.43 ± 6.17 % (52) vs Isolation = 6.81 ± 3.79 % (29) | Student’s *t*-test, 1.283, 79, *p* = 0.2032 |
| **Figure No.** | **Experiment** | **Mean ± SD (n)** | **Statistical test, t, df, *p*** |
| Figure 8C | %PPGNs *Gcg* only | Group = 54.78 ± 7.21 % (9) vs Isolation = 50.12 ± 20.44 % (13) | Student’s *t*-test, 0.6525, 20, *p* = 0.5215 |
|  | %PPGNs *Gcg* +*Htr1a* | Group = 15.54 ± 5.58 % (8) vs Isolation = 16.26 ± 15.81 % (13) | Student’s *t*-test, 0.1229, 19, *p* = 0.9034 |
|  | %PPGNs *Gcg* +*Htr2c* | Group = 15.01 ± 5.09 % (9) vs Isolation = 17.45 ± 8.73 % (13) | Student’s *t*-test, 0.7497, 20, *p* = 0.4622 |
|  | %PPGNs *Gcg* +*Htr1a +Htr2c* | Group = 11.65 ± 6.47 % (9) vs Isolation = 16.19 ± 9.52 % (13) | Student’s *t*-test, 1.242, 20, *p* = 0.2287 |
| Figure 8D | *Gcg* spots/ PPGN | Group = 19.51 ± 15.39 (9) vs Isolation = 32.36 ± 14.22 (12) | Student’s *t*-test, 1.979, 19, *p* = 0.0624 |
|  | *Htr1a* spots/ PPGN | Group = 9.71 ± 6.82 (9) vs Isolation = 4.06 ± 3.00 (12) | Mann-Whitney U, **p* = 0.0227 |
|  | *Htr2c* spots/ PPGN | Group = 9.73 ± 4.11 (9) vs Isolation = 13.84 ± 6.93 (13) | Student’s *t*-test, 1.587, 20, *p* = 0.1282 |

| **Figure No.** | **Experiment** | **Mean ± SD (n)** | **Statistical test, t, df, *p*** |
| --- | --- | --- | --- |
| Supp Figure 1A | Open Field- first 10’ distance traveled | Group = 6.13 ± 0.48 m (10) vs Isolation = 4.98 ± 0.77 m (10) | Student’s *t*-test, 4.040, 18, ****p* = 0.0008 |
|  | Open Field- first 10’ time in center | Group = 56.48 ± 16.76 s (10) vs Isolation = 62.24 ± 28.02 s (10) | Student’s *t*-test, 0.5575, 18, *p* = 0.5841 |
|  | Open Field- first 10’ time in corners | Group = 339.9 ± 59.6 s (10) vs Isolation = 338.5 ± 98.7 s (10) | Student’s *t*-test, 0.03806, 18, *p* = 0.9701 |
| Supp Figure 1B | Open Field- first 20’ distance traveled | Group = 11.37 ± 1.11 m (10) vs Isolation = 9.03 ± 1.69 m (10) | Student’s *t*-test, 3.662, 18, ***p* = 0.0018 |
|  | Open Field- first 20’ time in center | Group = 120.8 ± 29.2 s (10) vs Isolation = 137.2 ± 58.6 s (10) | Student’s *t*-test, 0.7895, 18, *p* = 0.4401 |
|  | Open Field- first 20’ time in corners | Group = 673.0 ± 130.8 s (10) vs Isolation = 662.8 ± 184.8 s (10) | Student’s *t*-test, 0.1418, 18, *p* = 0.8888 |
| Supp Figure 1C | Open Field- full 30’ distance traveled | Group = 16.31 ± 1.67 m (10) vs Isolation = 12.90 ± 2.53 m (10) | Student’s *t*-test, 3.558, 18, ***p* = 0.0022 |
|  | Open Field- full 30’ time in center | Group = 175.0 ± 43.3 s (10) vs Isolation = 204.3 ± 98.8 s (10) | Student’s *t*-test, 0.8603, 18, *p* = 0.4009 |
|  | Open Field- full 30’ time in corners | Group = 1013 ± 187 s (10) vs Isolation = 1007 ± 280 s (10) | Student’s *t*-test, 0.05449, 18, *p* = 0.9571 |

| **Figure No.** | **Experiment** | **Statistical Test, F stat** | **Multiple comparisons** |
| --- | --- | --- | --- |
| Supp Figure 3A | VO2 Dark phase | 2-w mixed RM ANOVA, Dark: Housing Condition F (1, 38) = 0.3359, *p* = 0.5656 |  |
|  | VO2 light phase | 2-w mixed RM ANOVA, Light: Housing Condition F (1, 38) = 0.5910, *p* = 0.4468 |  |
| Supp Figure 3B | Thermogenesis dark phase | 2-w mixed RM ANOVA, Dark: Housing Condition F (1, 38) = 0.3780, *p* = 0.5423 |  |
|  | Thermogenesis light phase | 2-w mixed RM ANOVA, Light: Housing Condition F (1, 38) = 0.1419, *p* = 0.7085 |  |
| Supp Figure 3C | Time budget | 2-w mixed RM ANOVA w/Geisser-Greenhouse’s epsilon, Housing Condition F(1.013, 20.55) = 73.79 | Sidak’s multiple comparisons, Group vs Isolation: Resting inside habitat, **p* = 0.0143; Long lounge (>60 s) **p* = 0.0213 |
